# Molecular genetic contributions to self-rated health

**DOI:** 10.1101/029504

**Authors:** Sarah E. Harris, Saskia P Hagenaars, Gail Davies, W David Hill, David CM Liewald, Stuart J Ritchie, Riccardo E Marioni, METASTROKE consortium, International Consortium for Blood Pressure, CHARGE consortium Aging and Longevity Group, CHARGE consortium Cognitive Group, Cathie LM Sudlow, Joanna M Wardlaw, Andrew M McIntosh, Catharine R Gale, Ian J Deary

**Author notes:** These authors contributed equally to the work (Saskia P Hagenaars, Gail Davies). These authors contributed equally to the work (Catharine R Gale, Ian J Deary). Corresponding author, Ian J. Deary Centre for Cognitive Ageing and Cognitive Epidemiology, Department of Psychology University of Edinburgh 7 George Square, Edinburgh, EH8 9JZ, Scotland, UK Telephone: *44, Fax: *44.

## Abstract

**Background:** Poorer self-rated health (SRH) predicts worse health outcomes, even when adjusted for objective measures of disease at time of rating. Twin studies indicate SRH has a heritability of up to 60% and that its genetic architecture may overlap with that of personality and cognition.

**Methods:** We carried out a genome-wide association study (GWAS) of SRH on 111 749 members of the UK Biobank sample. Univariate genome-wide complex trait analysis (GCTA)-GREML analyses were used to estimate the proportion of variance explained by all common autosomal SNPs for SRH. Linkage Disequilibrium (LD) score regression and polygenic risk scoring, two complementary methods, were used to investigate pleiotropy between SRH in UK Biobank and up to 21 health-related and personality and cognitive traits from published GWAS consortia.

**Results:** The GWAS identified 13 independent signals associated with SRH, including several in regions previously associated with diseases or disease-related traits. The strongest signal was on chromosome 2 (rs2360675, p = 1.77x10^−10^) close to KLF7, which has previously been associated with obesity and type 2 diabetes. A second strong peak was identified on chromosome 6 in the major histocompatibility region (rs76380179, p = 6.15x10^−10^). The proportion of variance in SRH that was explained by all common genetic variants was 13%. Polygenic scores for the following traits and disorders were associated with SRH: cognitive ability, education, neuroticism, BMI, longevity, ADHD, major depressive disorder, schizophrenia, lung function, blood pressure, coronary artery disease, large vessel disease stroke, and type 2 diabetes.

**Conclusions:** Individual differences in how people respond to a single item on SRH are partly explained by their genetic propensity to many common psychiatric and physical disorders and psychological traits.

**Key Messages:**

- Genetic variants associated with common diseases and psychological traits are associated with self-rated health.
- The SNP-based heritability of self-rated health is 0.13 (SE 0.006).
- There is pleiotropy between self-rated health and psychiatric and physical diseases and psychological traits.

## Introduction

There is considerable evidence that how individuals respond to one simple question asking them to evaluate their current state of health is a powerful predictor of future health outcomes. Poorer self-rated health (SRH) has been associated with increased mortality from all causes^1–4^ and from several specific causes including cardiovascular disease,^5–7^ diabetes,^8^ respiratory disease,^8^ cancer^8^ and infectious disease.^8^ Poorer SRH has also been linked in prospective studies to an increased risk of the onset of certain diseases, in particular, heart disease,^9–11^ cancer^11^ and type 2 diabetes,^12^ with a higher likelihood of incident admission to psychiatric hospital,^11^ and with increased incidence of cognitive or functional impairment.^13^ People with a greater burden of chronic disease are more likely to rate their health as poor or fair^14^ but, in general, adjustments for objective measures of disease, common risk factors, and health behaviours at the time that individuals rated their health, explains only a small part of the association between SRH and later morbidity or mortality.

Evidence for the heritability of SRH comes from several twin studies,^15–17^ which provide estimates of the percentage variance explained by genetic factors which range from ~20% to ~60%.^18,19^ Studies using molecular genetic methods also provide evidence for heritability: for instance, the genome-wide complex trait analysis (GCTA-GREML) method^20^ was used to estimate that common SNPs account for 18% of the variation in SRH (N = 4233). A multivariate twin study^22^ indicated appreciable genetic overlap between SRH and the phenotypically-correlated traits of optimism and self-rated mental health. However, there were also substantial genetic influences unique to SRH (see also^23^). In addition, Svedberg et al.^24^ showed that SRH and (measured) cognitive ability have a shared genetic basis using twin models; for older adults, genetic factors were entirely responsible for the phenotypic relation between SRH and cognitive ability. To date, studies have been insufficiently powered to detect variants from individual genes that relate to SRH.

Previous research suggests that perceptions of health are driven in part by psychological factors. There is evidence that people who are higher in the personality trait neuroticism—the tendency to experience negative emotions—are more likely to rate their health as being poor,^25–27^ and have a steeper decline in health ratings over time.^28^ Another psychological factor that has been linked with poorer SRH in cross-sectional surveys is lower cognitive ability. While there is some indication that poorer perception of health can be a risk factor for subsequent cognitive decline,^29^ longitudinal evidence suggests that having lower cognitive ability in youth increases the risk of poorer SRH decades later.^30^ Part of this link may be due to lower educational attainment—itself consistently linked with poorer SRH.^31–33^ It has been suggested that psychosocial resources may enable the highly educated to cope better with the negative effects of worsening health, and that this may in part explain why such individuals have better SRH.^33,35^

The aim of the present study was to add substantially to the understanding of the genetic mechanisms and genetic architecture of SRH. Using the large UK Biobank genotyped sample we conducted a genome-wide analysis of SRH, we estimated its SNP-based heritability, and we studied its pleiotropy with physical and mental health and with personality and cognitive traits.

## Methods

### Cohorts and measures

The UK Biobank is a health resource facilitating the study of the origins of a wide range of illnesses.^36^ Around 500 000 individuals aged between 37 and 73 years were recruited in the United Kingdom between 2006 and 2010. They underwent testing of cognitive abilities, physical and mental health examinations, completed questionnaires about lifestyle, sociod-emographic background and family medical history, and agreed to have their health followed longitudinally. For the present study, genome-wide genotyping data were available for 112 151 individuals (58 914 females, 53 237 males) aged 40 to 73 years (mean = 56.91 years, SD = 7.93). Figure 1 shows the participant selection for this study.

**Figure 1.**
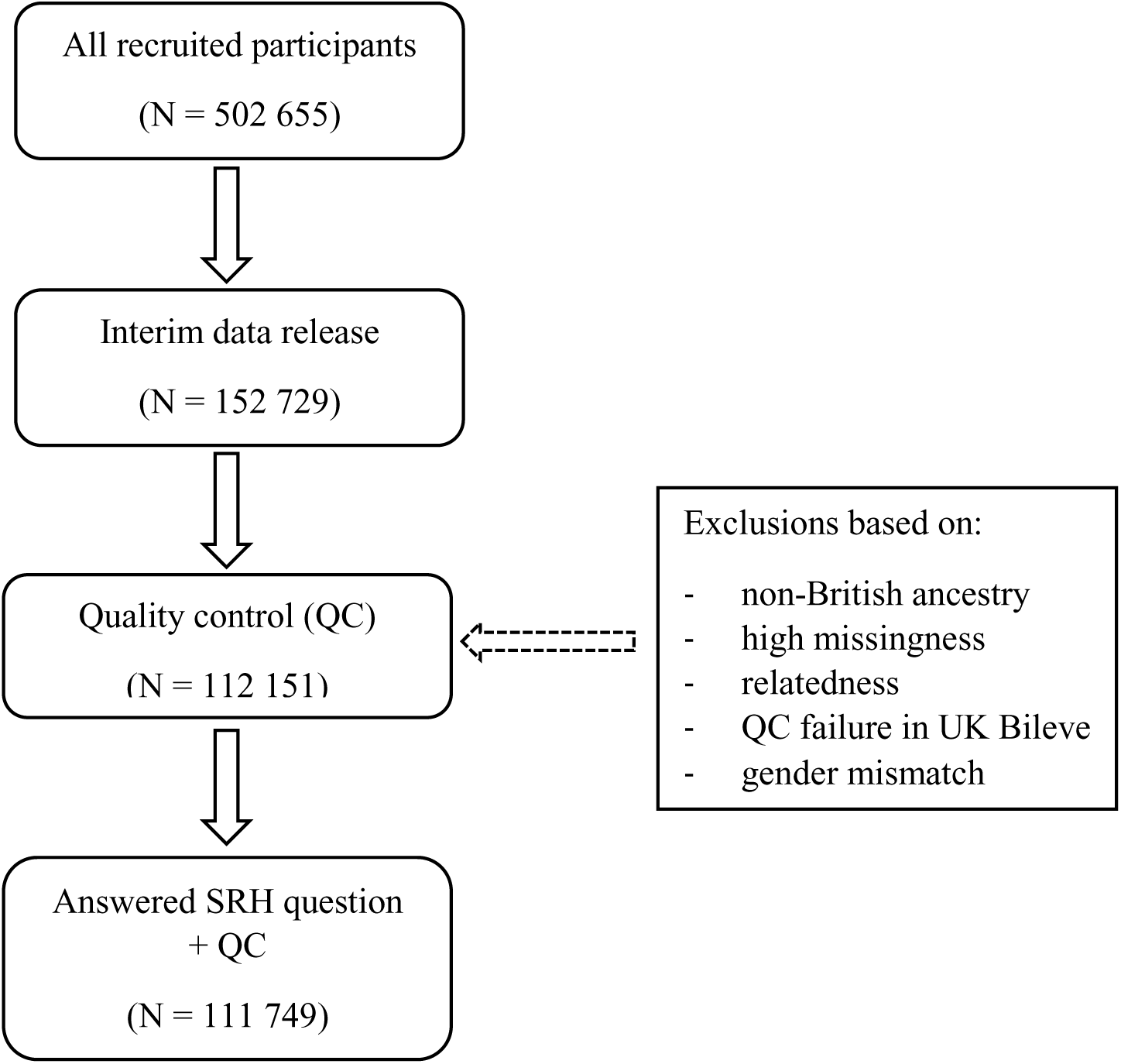
Flow diagram of participant selection

### Ethics

UK Biobank received ethical approval from the Research Ethics Committee (REC reference 11/NW/0382). This study has been completed under UK Biobank application 10279.

### Self-rated health

Participants were asked the question, “In general how would you rate your overall health?”. Possible answers were “Excellent/Good/Fair/Poor/Do not know/Prefer not to answer”. We created a four-category SRH variable indexing how each participant rated their health ranging from “excellent” to “poor”; excluding those that responded with “do not know” or “prefer not to answer”. For the phenotypic correlations, LD score regression and polygenic profile score analyses used in this study, a higher score for SRH indicates a better health rating.

### Neuroticism

Participants completed 12 questions of the Eysenck Personality Questionnaire-Revised Short Form (EPQ-R Short Form)^37,38^ neuroticism scale. Neuroticism refers to the relatively stable personality trait that assesses individual differences in the tendency to experience negative emotions. A summary score was derived to obtain a measure of neuroticism. The EPQ-R Short Form has been shown to correlate highly with other well-validated Neuroticism scales,^39^ and has shown a high genetic correlation (0.91) with psychological distress examined in a non-psychiatric population using the 30-item General Health Questionnaire.^40^

### Education

Education was measured by the question, “Which of the following qualifications do you have? (You can select more than one)”. Possible answers were: “College or University Degree/A levels or AS levels or equivalent/O levels or GCSE or equivalent/CSEs or equivalent/NVQ or HND or HNC or equivalent/Other professional qualifications e.g. nursing, teaching/None of the above/Prefer not to answer”. For the present study, a binary education variable was created to indicate whether or not a participant had a college or university-level degree; excluding those who responded with “prefer not to answer”. Previous studies have used similar binary variables as a ‘proxy-phenotype’ for cognitive ability.^41^

### Intelligence

Intelligence was measured by a thirteen item-test with a time limit of two minutes, completed by 36 035 individuals. Six items were verbal and seven numerical. An example of a verbal question is ‘Bud is to flower as child is to?’ (Possible answers: ‘Grow/Develop/Improve/Adult/Old/Do not know/Prefer not to answer’). An example of a numerical question is ‘If sixty is more than half of seventy-five, multiply twenty-three by three. If not subtract 15 from eighty-five. Is the answer?’ (Possible answers: ‘68/69/70/71/72/Do not know/Prefer not to answer’). The Intelligence score was the total score out of thirteen. The Cronbach α coefficient for the thirteen items was 0.62.

### Phenotypic Correlations

Phenotypic correlation coefficients were calculated between SRH, and neuroticism, education, intelligence and mortality in UK Biobank. Cox-proportional hazard ratios were calculated for all-cause mortality according to the SRH categories (Poor to Excellent).

### Genotyping and quality control

152 729 UK Biobank samples were genotyped using either the UK BiLEVE (N = 49 979) or the UK Biobank axiom array (N = 102 750). Array design, genotyping details and, quality control details can be found elsewhere.^42^ Genotyping was performed on 33 batches of ~ 4700 samples by Affymetrix. Initial quality control (QC) of the genotyping data was also performed by Affymetrix. Further details are available of the sample processing specific to the UK Biobank project (http://biobank.ctsu.ox.ac.uk/crystal/refer.cgi?id=155583) and the Axiom array (http://media.affymetrix.com/support/downloads/manuals/axiom_2_assay_auto_workflow_user_guide.pdf). Prior to the release of the UK Biobank genetic data a stringent QC protocol was applied, which was performed at the Wellcome Trust Centre for Human Genetics (WTCHG). Details of this process can be found here (http://biobank.ctsu.ox.ac.uk/crystal/refer.cgi?id=155580). Prior to the analyses described below, further quality control measures were applied. Individuals were removed sequentially based on non-British ancestry (within those who self-identified as being British, principal component analysis was used to remove outliers), high missingness, relatedness, QC failure in UK Bileve, and gender mismatch. A sample of 112 151 individuals remained for further analyses.

### Imputation

An imputed dataset was made available in which the UK Biobank interim release was imputed to a reference set combining the UK10K haplotype and 1000 Genomes Phase 3 reference panels. Further details can be found at the following URL: http://biobank.ctsu.ox.ac.uk/crystal/refer.cgi?id=157020. The association analyses were restricted to autosomal variants with a minor allele frequency greater than 0.1% and an imputation quality score of 0.1 or greater (N ~ 17.3m SNPs). An imputation quality score of 0.1 was chosen, rather than the more commonly used 0.3-0.5, due to the relatively large sample size of UK Biobank. An imputation score of 0.1 on 150 000 individuals corresponds to an effective sample size of 15 000 individuals.

### Curation of summary results from GWAS consortia on health-related variables

In order to conduct LD score regression and polygenic profile score analyses between the UK Biobank SRH and the genetic predisposition to psychiatric, physical and cognitive variables, we gathered 21 sets of summary results from international GWAS consortia of physical and psychiatric diseases and health-related traits and three sets of summary results from GWAS of the following UK Biobank variables: neuroticism, education and intelligence. Details of the health-related variables, the consortia’s websites, key references, and number of subjects included in each consortia’s GWAS are given in Supplementary Materials and Supplementary Table 1.

### Association analyses

The UK Biobank measure of SRH was adjusted for age, gender, assessment centre, genotyping batch, genotyping array, and 10 principal components for population stratification prior to the association analyses. The distribution of SRH was visually inspected and no exclusions were made; 111 749 individuals with both SRH and genotype information remained for further analyses.

SNPTEST v2.5.1^43^ was used to perform genotype-phenotype association analyses on the imputed dataset. SNPTEST v2.5.1 can be found at the following URL: https://mathgen.stats.ox.ac.uk/genetics_software/snptest/snptest.html#introduction. The ‘frequentist 1’ option was used to specify an additive model. Genotype dosage scores were used to account for genotype uncertainty.

The number of independent signals for the genotype-phenotype analyses was determined using LD clumping, using the 1000 genomes as a measure of LD between SNPs. First, SNPs with a genome-wide significant association with SRH (p < 5×10^−8^) were selected as index SNPs. Second, SNPs within 500kb and in LD of r^2^ > 0.1with the index SNP were included in the clump. SNPs from within this region were assigned to the clump if they had a P-value <1x10^−5^. In addition, conditional analyses were performed using SNPTEST v2.5.1^43^ for each genome-wide significant region with evidence of multiple signals, to identify potential secondary signals.

HLA classical allele imputation was performed using HLA genotype imputation with attribute bagging (HIBAG).^44^ HLA classical alleles were imputed in six genes (*HLA-A, HLA-B, HLA-C, HLA-DQA1, HLA-DQB1* and *HLA-DRB1*). Individuals with imputation score < 0.8 and alleles with frequency < 0.01 were removed prior to analysis. Linear regression correcting for age, gender, assessment centre, genotyping batch, genotyping array, imputation quality score and 10 principal components was performed to identify associations between HLA alleles and SRH.

MAGMA^45^ was used to perform gene-based association analyses. The results of the GWAS were used to derive the gene-based statistics. Genetic variants were assigned to genes based on their position according to the NCBI 37.3 build with no additional boundary placed around the genes; this resulted in a total of 18 116 genes being analysed. The European panel of the 1000 Genomes data (phase 1, release 3) was used as a reference panel to account for linkage disequilibrium. A genome-wide significance threshold for gene-based associations was calculated using the Bonferroni method (α= 0.05/18 116; P < 2.76 × 10^−6^).

### Functional annotation and gene expression

For the 13 independent genome-wide significant SNPs identified by LD clumping, evidence of expression quantitative trait loci (eQTL) and functional annotation were explored using publicly available online resources. The Genotype-Tissue Expression Portal (GTEx) (http://www.gtexportal.org) was used to identify eQTLs associated with the SNPs. Functional annotation was investigated using the Regulome DB database^46^ (http://www.regulomedb.org/). Regulome DB was used to identify regulatory DNA elements in non-coding and intergenic regions of the genome in normal cell lines and tissues.

### Estimation of SNP-based heritability

Univariate GCTA-GREML^20^ analyses were used to estimate the proportion of variance explained by all genotyped autosomal SNPs for SRH with a minor allele frequency > 0.01. A relatedness cut-off of 0.025 was used in the generation of the genetic relationship matrix.

### Genetic analyses: DEPICT

DEPICT^47^ was used to conduct three analyses; gene prioritisation, gene-set analysis, and tissue enrichment. The full GWAS output of SRH was clumped using PLINK to derive independent regions of the genome showing evidence of association. Next, DEPICT was used to determine if these independent regions overlapped with genes that share biological function by comparing the empirically-derived clumps with randomly-selected loci drawn from across the genome and matched for gene density. DEPICT tests the hypothesis that genes showing a true association with SRH will be involved in the same mechanisms that in turn contribute toward this phenotype. Clumping was performed using index SNPs of p < 1x10^−5^ with a 500kb boundary including SNPs in LD of r^2^ > 0.1.

### Stratified LD score regression

Stratified LD score regression was used in order to determine which regions of the genome are contributing towards variation in self-rated health. We follow the same data processing pipeline as Finucane et al.^48^ We first partitioned the full genome-wide data set into 24 functional annotations. An additional 500 bp boundary was placed around each of the regions captured by the annotations in order to avoid estimates being biased upwards by capturing enrichment from nearby SNPs, and a 100bp window was placed around chromatin immunoprecipitation and sequencing (ChIP-seq) peaks. This resulted in a baseline model consisting of a total of 52 overlapping functional annotations. The 24 main annotations included gene-sets from the digital genomic footprint (DGF ENCODE) and transcription factor binding sites (TFBS),^49,50^ DNase I hypersensitivity sites (DHS),^51^ the foetal gene-set being only those that were found within the foetal cell whereas the DHS grouping corresponding to all sites. Coding regions, 3’UTR, 5’UTR, promoter and intron^50,52^ gene-sets were used. Regions of the genome that have been evolutionarily conserved along the mammalian line^53,54^ were also included as were CTCF, promoter-flanking, transcribed, transcription start sites (TSS), strong enhancer and weak enhancer categories.^55^ Finally, cell type specific H3K4me1, H3K4me3, and H3K9ac data were taken from work performed on the Epigenomics Roadmap.^56^ An additional version of H3k27ac was also included^57^ as were clusters of enhancers that show a high level of activity (Super enhancers).^57^ This resulted in the 52 groupings that formed the baseline model.

In order to examine the role of specific tissue types, we then grouped the four histone marks (H3K4me1, H3K4me3, H3K9ac and H3K27ac) into 10 broad tissue types. These 10 groups correspond to histone marks found in the central nervous system (CNS), Immune and hematopoietic, adrenal/pancreas, cardiovascular, connective tissue, gastrointestinal, kidney, liver, skeletal muscle, and other.

The heritability Z-score for the SRH data set was 15.36 indicating that there was a sufficiently large polygenic signal for use with stratified LD score regression. The heritability of regions of the genome can be derived using stratified LD score regression.^48^ This heritability estimate can then be used to derive an enrichment metric, defined as the proportion of heritability the region captures over the proportion of SNPs that lie within the functional annotation, Pr(h^2^)/Pr(SNPs). Although using LD score regression of GWAS summary statistics that have undergone correction for population stratification can result in an attenuation of the heritability estimate derived, this will not bias the enrichment metric as regions of the genome will be influenced equally. LD scores were calculated from the European samples in the 1000 Genomes project (1000G) and only included the HapMap 3 SNPs with a minor allele frequency (MAF) of >0.05. False discovery rate (FDR)^58^ was applied to the full baseline model (52 tests) in order to control for the number of tests performed. For the tissue specific analysis, 10 tests were controlled for using FDR correction.

### Gene-set enrichment analysis

Gene-set enrichment analysis was performed using MAGMA.^45^ The full set of MsigDB canonical pathways^59^ was used to examine if any of the gene sets contained showed an association with self-rated health. Competitive testing was used to determine statistical significance, as this provides a means of correcting for the baseline level of association^45^ found in the self-rated health data set. The competitive test used in MAGMA is equivalent to comparing the mean association of the genes within each gene-set to the mean association of all the genes not contained within the gene-set. FDR correction was used to control for the number of tests performed.

Two methods have been used to compute genetic associations between health-related variables from GWAS consortia and SRH in UK Biobank: LD score regression and polygenic profile score analyses, both providing a different metric to examine pleiotropy between two traits. LD score regression was used to determine the degree of overlap in polygenic architecture between two traits by deriving genetic correlations. The polygenic profile score method was used to predict the phenotypic variance in SRH using summary data from GWASs of health-related variables to create polygenic profile scores in the UK Biobank sample. Both LD score regression and polygenic profile score analyses depend on traits being highly polygenic in nature, i.e. a large number of variants of small effect contributing toward phenotypic variation.^60,61^ LD score regression was performed between the 16 health related traits from GWAS consortia and three UK Biobank traits, while the polygenic profile score analyses were performed on the complete set of 21 health related traits from GWAS consortia as this method requires independent samples.

### LD score regression

LD score regression uses the information that for a given SNP, the effect size is a function of this particular SNP’s LD with other SNPs.^60,62^ Assuming a trait with a polygenic architecture, SNPs with high LD will have stronger effects on average than SNPs with low LD. LD score regression estimates the genetic effect on a trait by measuring the extent to which the observed effect sizes from a GWAS can be explained by LD. The covariance between the genetic effects in two traits can be indexed in a similar way, normalizing this genetic covariance by the heritability of the trait will estimate the genetic correlation between the two traits.

In the present study, LD score regression has been used to derive genetic correlations between summary statistics from 16 health related GWAS consortia and three UK Biobank GWA studies (Intelligence, Education and Neuroticism), and the UK Biobank SRH measure. We followed the data processing pipeline devised by Bulik-Sullivan et al.^60^ In order to ensure that the genetic correlation for the Alzheimer’s disease phenotype was not driven by a single locus or biased the fit of the regression model, a 500kb region centred on the *APOE* locus was removed and this phenotype was re-run. This additional model is referred to in the Tables and Figures as ‘Alzheimer’s disease (500kb)’.

### Polygenic profile score analyses

The UK Biobank genotyping data required recoding from numeric (1, 2) allele coding to standard ACGT format prior to being used in polygenic profile scoring analyses. This was achieved using a bespoke programme developed by one of the present authors (DCML), details of which are provided in the Supplementary Materials.

PRSice^63^ was used to create polygenic profile scores from 21 health-related phenotypes of published GWAS in all genotyped participants (Supplementary Table 1). SNPs with a minor allele frequency < 0.01, as well as strand-ambiguous SNPs were removed prior to creating the scores. Clumping was used to obtain SNPs in linkage equilibrium with an r^2^ < 0.25 within a 200bp window. The scores were calculated as the sum of alleles associated with the phenotype of interest across many genetic loci, weighted by their effect sizes estimated from the GWAS summary statistics. The conventional approach was used to create polygenic profile scores that included variants according to the significance of their association with the phenotype, exceeding five predefined p-value thresholds of 0.01, 0.05, 0.1, 0.5 and all SNPs. Throughout the paper, the most predictive threshold will be presented in the main tables; the full results, including all five thresholds, can be found in Supplementary Table 10.

Regression models were used to examine the associations between the 21 polygenic profiles and SRH, adjusting for age at measurement, sex, genotyping batch and array, assessment centre, and the first ten genetic principal components to adjust for population stratification.

All polygenic profile score association analyses were performed in R,^64^ and the obtained p-values from each test were corrected for multiple testing using the False Discovery Rate (FDR) method.^58^ Sensitivity analyses were performed in order to test whether the results are driven by individuals with a given illness. This was done by removing individuals with a self-reported clinical diagnosis of coronary artery disease (N = 5300), type 2 diabetes (N = 5800) and hypertension (N = 26 912) from the relevant analyses. More details can be found in the Supplementary Materials. Multivariate regression has been performed including all FDR significant polygenic profile scores and the covariates described earlier.

### Mendelian Randomisation

To begin to address causality, a Mendelian Randomisation approach was used to investigate if genetically determined BMI was associated with SRH in UK Biobank. A polygenic risk score (PGRS) for BMI was created, using PRSice,^63^ for each UK Biobank subject using the SNPs associated with BMI at a genome-wide association significance level in the GIANT consortium.^65^ 90 of the 97 SNPs were available and not strand-ambiguous in the UK Biobank data set (Supplementary Table 2). Linear regressions correcting for age, gender, assessment centre, genotyping batch, genotyping array, and 10 principal components were used to examine the associations between BMI and BMI PGRS, and between SRH and BMI PGRS. The PGRS was then used as an instrumental variable in a two stage least squares regression (performed in R using the sem package) correcting for age, gender, assessment centre, genotyping batch, genotyping array, and 10 principal components to test the potential causal role of BMI on SRH.

## Results

### Phenotypic correlations

Within UK Biobank, 111 749 individuals with genotype data completed the question ‘How would you rate your overall health?’ Their mean (SD) score for SRH was 2.14 (0.73). SRH showed a negative correlation with the measure of neuroticism (r = -0.25, p < 0.0001), indicating that individuals who rate their health as worse had higher levels of neuroticism. Correlations were also found for the UK Biobank measures of intelligence and education (r = 0.146 and 0.110, p < 0.0001), indicating that individuals with higher levels of intelligence or education are more likely to rate their health as better. Cox proportional hazard models for all-cause mortality adjusted for age and sex, indicated that, compared to people with excellent SRH, the risk of dying in those with good, fair or poor SRH is 1.37 (1.17, 1.62), 2.51 (2.12, 2.97), 6.95 (5.79, 8.36) respectively.

### Genome-wide association study

A total of 109 SNPs from 12 genomic regions were associated with SRH (Figure 2, Figure 3, Table 1 and Supplementary Table 3). Thirteen independent signals were identified by clumping. Conditional analyses did not identify any secondary signals (Supplementary Figure 1). The strongest signal was on chromosome 2 and included the gene encoding Kruppel-Like Factor 7 (*KLF7*). Variants in this gene have previously been associated with obesity^66^ and type 2 diabetes.^67^ A second strong peak was identified on chromosome 6. Clumping identified two SNPs within the major histocompatibility complex region (MHC). Conditional analyses indicate that they are not independent. The MHC consists of a large number of genes that encode a group of cell surface molecules, which have important roles in the immune system. HLA allele analysis indicated that the allele with the strongest association with SRH is HLA-DQB1*03.02 (standardised beta = 0.029) (Supplementary Tables 4-9). Two independent signals were identified on chromosome 3, one of which was within Bassoon Presynaptic Cytomatrix Protein (*BSN*), a gene that encodes a scaffold protein expressed in the brain, is involved with neurotransmitter release and was previously associated with Crohn’s disease.^68^ A single SNP in Sterile Alpha Motif Domain Containing 12 (*SAMD12*) on chromosome 8 was associated with SRH. This region has previously been linked to diastolic blood pressure.^69^ A single SNP in Transcription Factor 4 (*TCF4*, chromosome 18), believed to be important in nervous system development and previously associated with neurodevelopmental disorders and psychiatric diseases was also associated with SRH.^70^ Single SNPs were also identified in SEC24 Family Member C (*SEC24C*) involved in vesicle trafficking and Shisa Family Member 9 (*SHISA9*) a regulator of short-term plasticity in the dentate gyrus, on chromosomes 10 and 16 respectively.^71,72^

**Figure 2.**
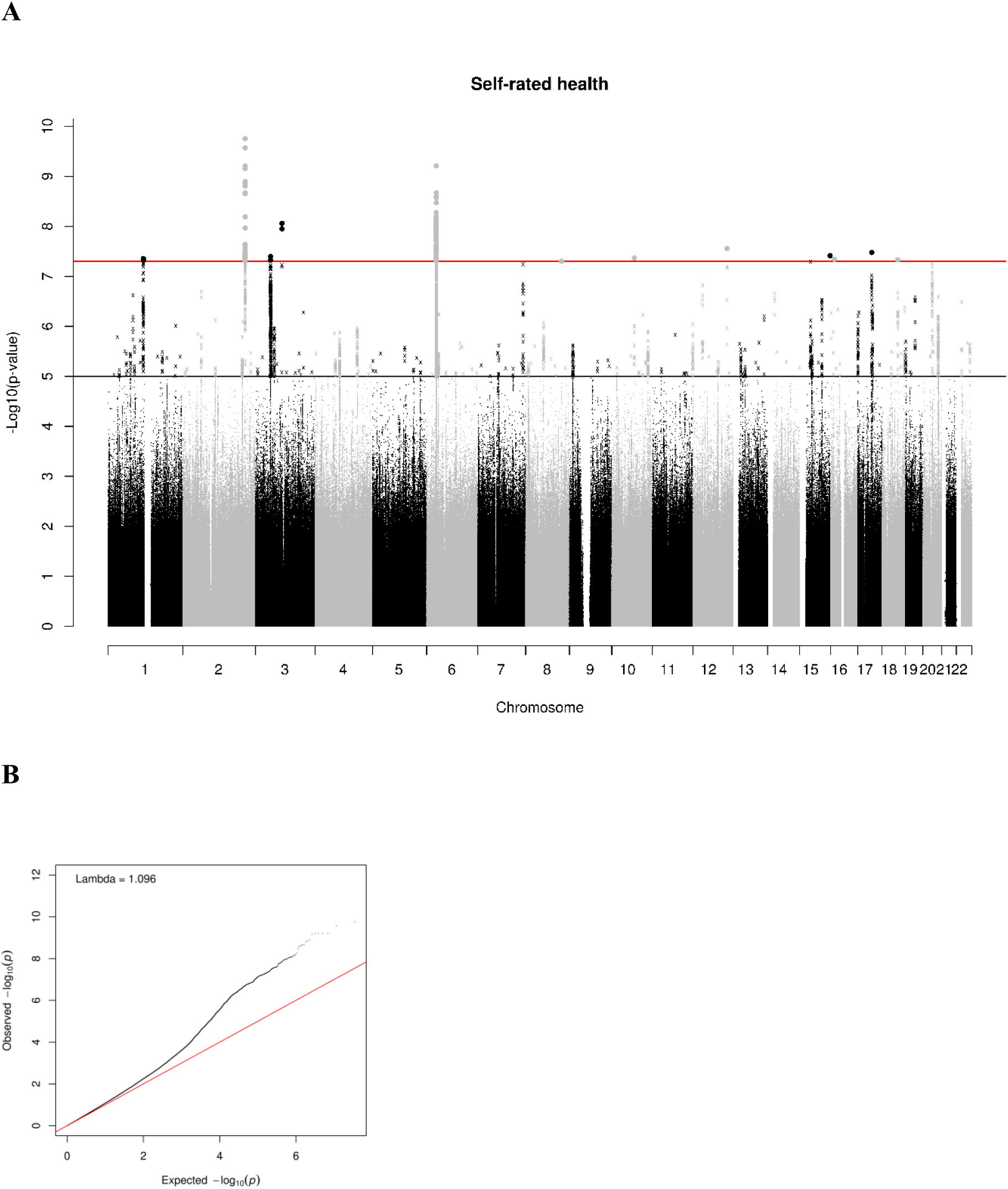
(A) Manhattan and (B) Q-Q plot of P-values of the SNP-based association analysis. The red line indicates the threshold for genome-wide significance (P<5 x 10-8); the grey line indicates the threshold for suggestive significance (P<1 x 10-5).

**Figure 3.**
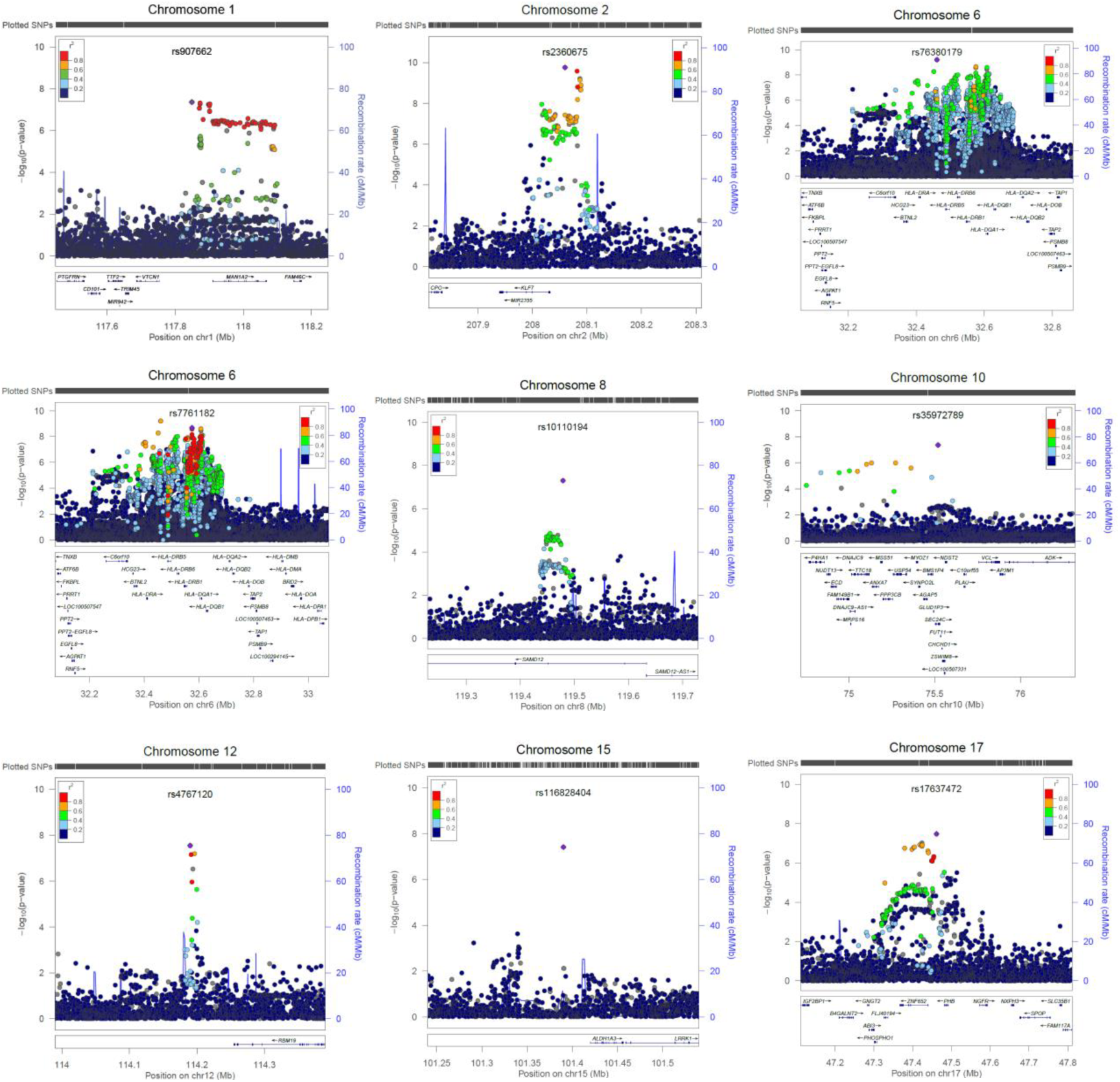
Regional association plots of genomic regions that demonstrated genome-wide significance (P<5 x 10-8) in the SNP-based association analyses for self-rated health. The circles represent individual SNPs, with the colour indicating pairwise linkage disequilibrium (LD) to the SNP indicated by the purple diamond (calculated from 1000 Genomes Nov 2014 EUR). The purple diamond indicates the most significant SNP for which LD information was available in the 1000G reference sample. The solid blue line indicates the recombination rate and -log10 P-values are shown on the y-axis.

**Table 1.** Independent genome-wide significant SNP-based associations for self-rated health.

| SNP | Chr | Position | A 1 | A 2 | N | p | $\beta$ | MAF | INFO | Genes in region | Previous gene associations |
| --- | --- | --- | --- | --- | --- | --- | --- | --- | --- | --- | --- |
| rs2360675 | 2 | 208059640 | A | C | 111749 | $1.77 \times 10^{-10}$ | -0.027 | 0.44 | 0.99 | *KLF7* | obesity<sup>66</sup> and type 2 diabetes<sup>67</sup> |
| rs76380179 | 6 | 32460750 | T | C | 111749 | $6.15 \times 10^{-10}$ | 0.038 | 0.17 | 0.82 | HLA Class II | auto immune diseases<sup>80</sup> |
| rs7761182 | 6 | 32575099 | T | G | 111749 | $2.13 \times 10^{-9}$ | 0.030 | 0.22 | 0.97 | HLA Class II | auto immune diseases<sup>80</sup> |
| rs528886319 | 3 | 87511141 | A | G | 111749 | $1.12 \times 10^{-8}$ | -0.26 | 0.0029 | 0.78 | NA | NA |
| rs4767120 | 12 | 114189961 | G | A | 111749 | $2.79 \times 10^{-8}$ | -0.024 | 0.40 | 0.98 | AK096932/BC007399 | NA |
| rs17637472 | 17 | 47461433 | A | G | 111749 | $3.32 \times 10^{-8}$ | 0.024 | 0.40 | 0.98 | NA | NA |
| rs116828404 | 15 | 101390343 | A | G | 111749 | $3.87 \times 10^{-8}$ | 0.45 | 0.0012 | 0.55 | LOC145757 | NA |
| rs147141470 | 3 | 49638084 | A | AAA<br>ATT | 111749 | $4.01 \times 10^{-8}$ | -0.026 | 0.29 | 0.98 | *BSN* | Crohn's disease<sup>68</sup> |
| rs35972789 | 10 | 75519691 | A | C | 111749 | $4.30 \times 10^{-8}$ | -0.061 | 0.038 | 1 | *SEC24C* | NA |
| rs907662 | 1 | 117848822 | A | G | 111749 | $4.41 \times 10^{-8}$ | -0.027 | 0.24 | 0.99 | NA | NA |
| rs555785137 | 16 | 13015115 | G | A | 111749 | $4.56 \times 10^{-8}$ | -0.45 | 0.0012 | 0.58 | *SHISA9* | NA |
| rs2924322 | 18 | 53244414 | T | A | 111749 | $4.68 \times 10^{-8}$ | 0.039 | 0.12 | 0.84 | *TCF4* | neurodevelopmental disorders and psychiatric diseases<sup>70</sup> |
| rs10110194 | 8 | 119479124 | C | G | 111749 | $4.98 \times 10^{-8}$ | 0.025 | 0.35 | 0.95 | *SAMD12* | diastolic blood pressure<sup>69</sup> |

### Gene-based analyses (MAGMA)

The gene-based analysis identified 36 genes across 11 genomic regions associated with SRH (Supplementary Table 10). The most significantly associated gene was *BSN*. Eighteen other genes in this gene dense region of chromosome 3 are also included in the list. This same region previously showed suggestive significance with general cognitive function.^73^ Four major histocompatibility complex genes (*HLA-DQA1, HLA-DQB1, HLA-DRB1* and *HLA-DRB5*) were also associated with SRH. Other genes of potential interest include: neurexin 1 (*NRXN1*, chromosome 2), a synaptic adhesion molecule previously associated with neurodevelopmental disorders;^74^ autism susceptibility candidate 2 (*AUTS2*, chromosome 7), previously associated with neurodevelopmental disorders and cancer;^75^ Zinc Finger Protein 652 (*ZNF652*, chromosome 17) a zinc finger protein previously associated with blood pressure;^76^ and Additional Sex Combs Like Transcriptional Regulator 3 (*ASXL3*, chromosome 18), previously associated with cancer.^77^

### GCTA-GREML analysis of SNP-based heritability

The proportion of variance in SRH that was explained by all common genetic variants was 13% (GCTA-GREML estimate 0.13, SE 0.006).

### Functional annotation and gene expression

Using the GTEx database (http://www.broadinstitute.org/gtex/), three cis-eQTL associations were identified for the 13 independent genome-wide significant SNPs (Supplementary Table 11). rs907662 on chromosome 1 potentially regulates Mannosidase, Alpha, Class 1A, Member 2 (*MAN1A2*), previously identified as being differentially expressed in type 2 diabetes patients compared to normal controls.^78^ rs76380179 and rs7761182 on chromosome 6 potentially regulate a number of major histocompatibility genes. There was evidence of regulatory elements associated with all nine of the independent genome-wide significant SNPs included in the Regulome DB database. (http://www.regulomedb.org/) (Supplementary Table 11).

### Gene prioritisation, gene set analysis and tissue enrichment

The gene prioritisation analysis, gene set analysis and the tissue enrichment, performed in DEPICT, provided no evidence of association for any of the gene sets or tissue types considered. Full results for these analyses can be found in Supplementary Tables 12, 13 and 14.

### Stratified LD score regression

The baseline model yielded eight significantly enriched functional annotations (Supplementary Figure 2). This indicates that the SNPs within these annotation categories explain a greater proportion of the variance for SRH than would be expected based on the proportion of the total number of SNPs used for this analysis. SNPs in evolutionarily conserved regions showed enrichment as they account for only 2.6% of the total number of SNPs but explain 39% of the heritability (enrichment metric = 15, SE = 2.7, P = 1.27x10-7. Enrichment was also found in super enhancers, which contained 16.8% of the SNPs collectively explaining 27.9% of the heritability (enrichment metric = 1.66, SE = 0.16, P = 5.42x10-5). SNPs within 500 bp of DNase hypersensitivity sites (DHS) also showed significant enrichment, accounting for 50% of the total SNPs and explaining total of 84.2% of the heritability (enrichment metric = 1.69, SE 0.197, P = 0.00049). A number of histone markers were also enriched including the H3K27ac mark, (enrichment metric = 1.37, SE 0.129, P = 0.0041), the H3K4me1 mark (enrichment metric = 1.71, SE 0.25, P = 0.00428), and SNPs within 500 bp of the H3K9ac mark (enrichment metric = 1.84, SE 0.308, P = 0.0063). The results of the cell type specific histone marks are shown in Supplementary Figure 3. Histone marks that are present in the central nervous system were significantly enriched. These SNPs contain 14.8% of the total number of SNPs which collectively explained 42.8% of the heritability (enrichment metric = 2.88, SE = 0.360, P = 1.68x10-7).

### Gene-set enrichment analysis

None of the gene-sets examined showed statistical significance once correction for multiple comparisons had been applied. Supplementary Table 15 shows the full output of the gene-set enrichment analysis.

To test for pleiotropy between SRH and health-related, personality and cognitive traits, we present LD score regression and polygenic profile analyses. For the purpose of these two analyses, a higher score for SRH indicates a better health rating.

### LD score regression

LD score regression was performed to obtain genetic correlations between SRH in UK Biobank and the summary results of the 16 GWAS consortia and three UK Biobank traits (Neuroticism, Education and Intelligence) (Figure 4, Supplementary Table 16). Better SRH showed positive genetic correlations with intelligence, education, longevity, anorexia nervosa, and forced expiratory volume in one second (FEV_1_) (rg = 0.11 to 0.59). Negative genetic correlations were found between better SRH and neuroticism, BMI, ADHD, major depressive disorder, schizophrenia, systolic and diastolic blood pressure, coronary artery disease, ischaemic stroke, and type 2 diabetes (rg = -0.16 to -0.46).

**Figure 4.**
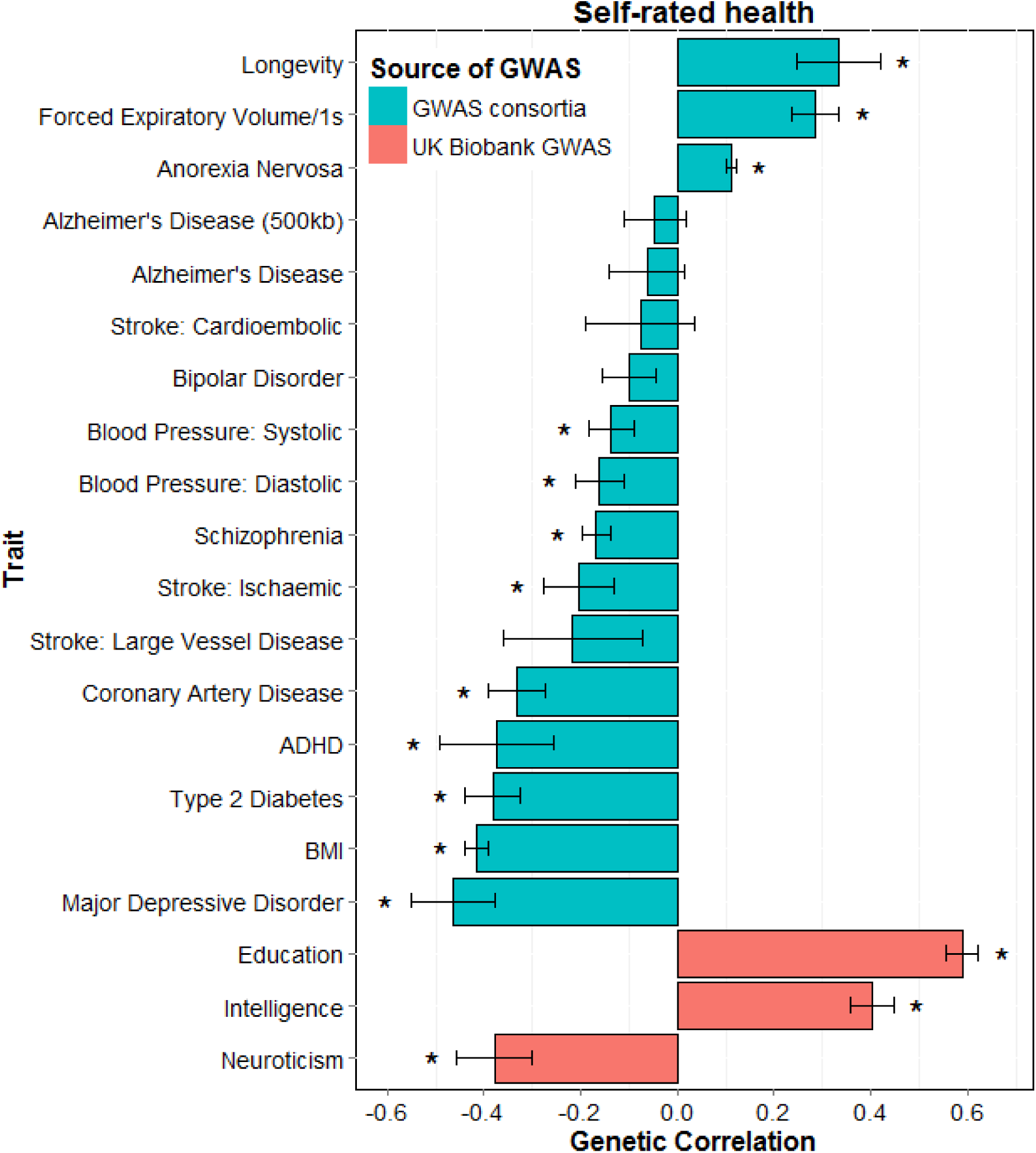
Barplot of genetic correlations calculated using LD score regression between self-rated health in UK Biobank (N = 111 749) and 15 health-related measures from GWAS consortia and three from UK Biobank (Intelligence, Education and Neuroticism). Self-rated health is scored such that higher scores indicate a better health rating. *, FDR-corrected p < 0.0061.

### Polygenic profile analyses

The results of the polygenic risk score analyses are shown in Table 2, using the most predictive threshold for each trait. The numbers of SNPs included in each polygenic threshold score for each of the 21 health-related traits are shown in Supplementary Table 17. Higher polygenic profile scores for years of education, general- and childhood cognitive ability, longevity, and FEV1 were associated with higher levels of SRH (standardised β between 0.01 and 0.06). Higher polygenic profile scores for neuroticism, BMI, ADHD, major depressive disorder, schizophrenia, diastolic and systolic blood pressure, coronary artery disease, large vessel disease stroke, and type 2 diabetes were associated with lower levels of SRH (standardised β between -0.07 and -0.009). The results showed very little change when individuals with self-reported clinical diagnoses of cardiovascular disease (N = 5300), diabetes (N = 5800), and hypertension (N = 26 912) were removed from the corresponding analyses (coronary artery disease, type 2 diabetes and systolic blood pressure). The results including all five thresholds can be found in Supplementary Table 18.

**Table 2.** Associations between polygenic profiles of health related traits created from GWAS consortia summary data, and UK Biobank self-rated health controlling for age, sex, assessment centre, genotyping batch and array, and ten principal components for population structure (N = 111 749). FDR-corrected statistically significant values (P < 0.0324) are shown in bold. Self-rated health is scored such that higher scores indicate better health ratings. The associations between the polygenic profile scores with the largest effect size (thresh) and self-rated health are presented. Thresh; the p-value threshold with the largest effect size.

| **Trait** | **thresh** | **$\beta$** | **p** |
| --- | --- | --- | --- |
| Years of education | 0.5 | 0.058 | **$6.06 \times 10^{-80}$** |
| Childhood cognitive ability | 0.5 | 0.025 | **$1.02 \times 10^{-16}$** |
| General cognitive function | 1 | 0.037 | **$1.29 \times 10^{-30}$** |
| Neuroticism | 0.5 | -0.017 | **$1.89 \times 10^{-8}$** |
| BMI | 0.5 | -0.067 | **$8.20 \times 10^{-108}$** |
| Longevity | 0.05 | 0.011 | **$5.49 \times 10^{-4}$** |
| ADHD | 0.1 | -0.009 | **$2.13 \times 10^{-3}$** |
| Alzheimer's Disease | 1 | -0.003 | $3.50 \times 10^{-1}$ |
| Anorexia Nervosa | 0.01 | 0.007 | $3.04 \times 10^{-2}$ |
| Bipolar Disorder | 0.01 | -0.005 | $9.59 \times 10^{-2}$ |
| Major Depressive Disorder | 1 | -0.017 | **$3.44 \times 10^{-8}$** |
| Schizophrenia | 1 | -0.028 | **$2.14 \times 10^{-19}$** |
| FEV<sub>1</sub> | 1 | 0.020 | **$7.71 \times 10^{-12}$** |
| Blood Pressure: Diastolic | 0.5 | -0.012 | **$4.77 \times 10^{-5}$** |
| Blood Pressure: Systolic | 0.1 | -0.015 | **$9.05 \times 10^{-7a}$** |
| Coronary Artery Disease | 0.5 | -0.025 | **$7.19 \times 10^{-17b}$** |
| Stroke: Ischaemic | 0.05 | -0.007 | $3.24 \times 10^{-2}$ |
| Stroke: Cardioembolic | 0.05 | -0.006 | $4.11 \times 10^{-2}$ |
| Stroke: Large Vessel Disease | 0.05 | -0.010 | **$1.19 \times 10^{-3}$** |
| Stroke: Small Vessel Disease | 0.1 | -0.006 | $5.74 \times 10^{-2}$ |
| Type 2 Diabetes | 1 | -0.032 | **$1.88 \times 10^{-24d}$** |

<sup>a</sup> Excluding individuals with hypertension $\beta = -0.010$ , $p = 0.005$ ; <sup>b</sup> excluding individuals with cardiovascular disease $\beta = -0.018$ , $p = 8.4 \times 10^{-9}$ ; <sup>c</sup> excluding individuals with diabetes $\beta = -0.021$ , $p = 4.08 \times 10^{-11}$ . FEV<sub>1</sub> = forced expiratory volume in one second.

A multivariate regression model was run including 14 of 15 significant polygenic profile scores (years of education, childhood cognitive ability, general cognitive function, neuroticism, BMI, longevity, ADHD, major depressive disorder, schizophrenia, FEV_1_, systolic blood pressure, coronary artery disease, large vessel disease stroke, and type 2 diabetes) alongside the same covariates as described previously. Due to the high phenotypic correlation between systolic and diastolic blood pressure, only systolic blood pressure was included in the model. This tested the extent to which including all significant polygenic profile scores in a multivariate model would improve the prediction of SRH and discover which polygenic scores contributed independently. This was done by subtracting the r^2^ value of the model only including the covariates from the model with both covariates and polygenic profile scores. All polygenic profile scores remained significant, after FDR correction (p < 0.032), in this multivariate model, and together accounted for 1.03% of the variance in SRH (Table 3).

In UK Biobank BMI was negatively correlated with SRH (Spearman’s rank correlation = - 0.26). A 1 SD increase in BMI polygenic risk score, based on 90 SNPs, was associated with a 0.12 SD increase in BMI. A 1 SD increase in BMI polygenic risk score was associated with a 0.026 SD decrease in SRH. A genetically determined increase in BMI of 0.58kg/m^2^ was associated with a 0.45 SD decrease in SRH.

**Table 3.**
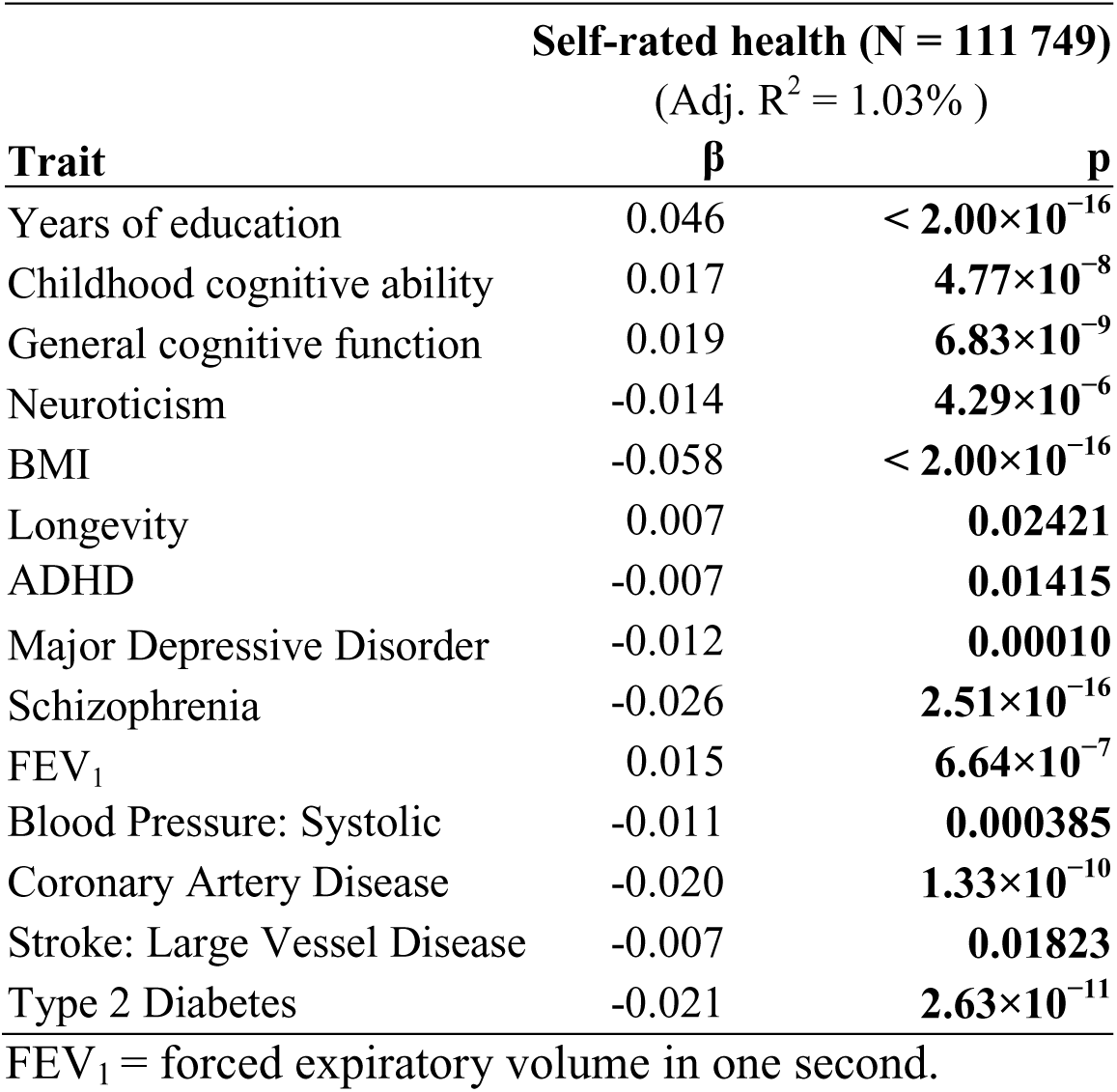
Multivariate models predicting self-rated health, including all significant polygenic profile scores together with covariates (age, sex, assessment centre, genotyping batch and array, and ten genetic principal components for population structure; covariate values not shown here). Self-rated health is scored such that higher scores indicate better health ratings. Adjusted R^2^ values refer to the polygenic profile scores only (excluding variance explained by the covariates). Statistically significant p-values (after FDR correction; threshold: P < 0.0243) are shown in bold.

| **Self-rated health (N = 111 749)** |  |  |
| --- | --- | --- |
| (Adj. $R^2 = 1.03\%$ ) | | |
| **Trait** | **$\beta$** | **p** |
| Years of education | 0.046 | **$< 2.00 \times 10^{-16}$** |
| Childhood cognitive ability | 0.017 | **$4.77 \times 10^{-8}$** |
| General cognitive function | 0.019 | **$6.83 \times 10^{-9}$** |
| Neuroticism | -0.014 | **$4.29 \times 10^{-6}$** |
| BMI | -0.058 | **$< 2.00 \times 10^{-16}$** |
| Longevity | 0.007 | **0.02421** |
| ADHD | -0.007 | **0.01415** |
| Major Depressive Disorder | -0.012 | **0.00010** |
| Schizophrenia | -0.026 | **$2.51 \times 10^{-16}$** |
| FEV<sub>1</sub> | 0.015 | **$6.64 \times 10^{-7}$** |
| Blood Pressure: Systolic | -0.011 | **0.000385** |
| Coronary Artery Disease | -0.020 | **$1.33 \times 10^{-10}$** |
| Stroke: Large Vessel Disease | -0.007 | **0.01823** |
| Type 2 Diabetes | -0.021 | **$2.63 \times 10^{-11}$** |

FEV<sub>1</sub> = forced expiratory volume in one second.

## Discussion

In the present and other studies, a single-item of SRH is associated with mortality. Such SRH items are widely and successfully used in health research. Given their predictive validity, it is of interest to discover the causes of people’s differences in SRH. Here, in analyses of the large UK Biobank sample together with results from many GWAS consortia, we discovered many new genome-wide significant genetic variants associated with SRH. A robust estimate of the SNP-based heritability of SRH in the UK was provided (13%), which is close to that previously reported (18%) using this method in the US based Health and Retirement Survey.^21^ Extensive pleiotropy was found between SRH and many physical and psychiatric disorders and health-related, cognitive and personality traits, indicating that, to a significant degree, the same genetic variants may influence these traits and SRH. This provides comprehensive new findings on the overlap between how individuals rate their health on a four-point scale and the genetic contributions to intelligence, personality, cardiovascular diseases, and many psychiatric and physical disorders and traits.

The present study identified novel genes/loci associated with individual differences in SRH. These include genes previously associated with diabetes (*KLF7, MAN1A2*),^67,78^ neurodevelopmental disorders (*TC4F, NRXN1, AUTS2*),^70,74,75^ autoimmune diseases (BSN),^68^ blood pressure (*SAMD12*, ZNF652)^69,76^ and cancer (*ASXL3*, AUTS2),^77,75^ although as yet not at a gold standard genome-wide significance level. These results indicate that genes previously associated with objectively measured diseases are also associated with SRH, perhaps indicating that people’s perception of their health does truly reflect their state of health. The MHC on chromosome 6 was also shown to be associated with SRH. The MHC is vital for the correct functioning of the immune system and therefore genetic variants in this region can have major health implications, for example *HLA-DQA1* which was associated with SRH in our gene-based analysis, has previously been associated with coeliac disease.^79^ HLA allele analysis indicated that the HLA-DQB1*03.02 was most strongly associated with SRH. This allele has previously been associated with several autoimmune diseases including type I diabetes and multiple sclerosis.^80,81^ Several genes were identified using the gene-based analysis in regions that did not contain any genome-wide significant SNPs indicating the importance of gene-based tests. None of our genome-wide significant SNPs were in regions that approached significance in a previous much smaller GWAS of SRH. ^19^

Sensitivity analyses showed that the polygenic profile score analyses for systolic blood pressure, coronary artery disease, and type 2 diabetes were not confounded by individuals with the associated disease (hypertension, cardiovascular disease and diabetes mellitus). This indicates that even in self-rated healthy individuals, a higher polygenic profile score for systolic blood pressure, coronary artery disease and type 2 diabetes is associated with lower health ratings.

The results of the present study indicate that genetic variants associated with better SRH are associated with a lower genetic risk of neuroticism, but a higher genetic risk of anorexia nervosa. From a previously published positive phenotypic association between anorexia nervosa and neuroticism,^82^ and our finding that individuals who rate their health lower had higher levels of neuroticism, one might have expected that high polygenic risk of anorexia nervosa would be associated with lower SRH. However, the polygenic profile score for anorexia nervosa could be seen on a spectrum, where individuals on the lower end of the spectrum might be more conscious about their eating behaviour and health, leading to better SRH, without exceeding the threshold for a clinical diagnosis of anorexia nervosa. The summary results of the GWAS for anorexia nervosa used for both LD score regression and polygenic profile analyses were based on 2907 cases and almost 15 000 controls.^83^ It is possible that this GWAS is picking some degree of predisposition to healthy behaviour. Another explanation for this finding is that individuals with anorexia nervosa potentially have a discrepancy between their SRH and their actual health, due to the body image distortion of individuals with anorexia nervosa. This study was unable to test this hypothesis.

This study shows that the SRH measure, consisting of only one question, is able to reflect the genetic variants of traits and disorders, such as intelligence, personality, cardio-metabolic disease and psychiatric disorders, associated with actual health. Genetic variants associated with higher levels of intelligence and lower levels of cardio metabolic diseases are associated with better health ratings. This supports the theoretical construct of bodily system integrity, a latent trait indicating individual differences in encountering health and cognitive challenges from the environment.^84^ Individuals with better system integrity are likely to have higher levels of intelligence, fewer diseases, a better overall health and greater longevity. A test of the hypothesis is whether there are genetic associations between any cognitive-related trait from youth and later health. Here, the clearest evidence for this is the large genetic correlation between education—which, for most people, is completed in youth—and self-rated health in older age.

The strongest association found in this study is between SRH and the polygenic profile score for BMI, accounting for 0.45% of the variance. Mendelian Randomisation indicated that higher genetically determined BMI leads to lower SRH, although we cannot exclude a causal effect in the other direction. When combining the polygenic liabilities for multiple traits and disorders in a multivariate model, the polygenic liabilities together double the amount of variance to 1%. This implies that SRH may be affected by risk alleles unique to each trait and disorder.

A strength of this study is the large sample size of UK Biobank, permitting powerful and robust tests of pleiotropy between SRH and many health related traits. Other strengths include that all individuals were of White British ancestry, minimising population stratification. Genotyping and quality control has been performed in a consistent way across the whole sample. The use of summary data from many international GWAS consortia allowed a detailed examination of pleiotropy between SRH and a wide range of health-related traits, showing many novel estimates of genetic correlations between traits.

The present study has some limitations; several of the genome-wide significant hits are single SNPs rather than in a peak. These SNPs should be treated with particular caution until they are replicated in an independent sample, which is currently not available. The summary data from the GWAS studies curated to perform LD score regression and create polygenic profile scores often originated from consortia studies, which involve meta-analyses across datasets with substantial heterogeneity in sample size, genome-wide imputation quality, and measurement of the traits. For the polygenic profile analyses we might have overestimated the effects because of possible overlap of individuals in UK Biobank sample and some of the cohorts within some of the GWAS consortia. We were unable to quantify the exact overlap, but the number of overlapping individuals is probably small and we judge that this will have a minor effect on the results. Because the analyses were restricted to individuals of White British ancestry, we are unable to generalize the results beyond that group. Therefore, these analyses should be replicated in large samples of individuals with different backgrounds. For the majority of traits we were unable to distinguish between type I pleiotropy (a single locus directly influencing multiple phenotypes) and type II pleiotropy (a single locus influencing a cascade of events). The findings from this study are not necessarily transferable to other populations as SRH is influenced by social and cultural components.

## Summary

Measuring people’s overall health is difficult, because the state of the body and mind can be disrupted in many ways, and people’s perceptions of the same objective bodily state can differ. Notwithstanding this complexity, the response to a single subjective question about whether a person is in good or poor health has proved valid and useful in health research. The present study has been able to identify many genetic contributions to SRH, confirming the complexity of the contributions to the phenotype, and also its partial foundations in genetic differences. The mechanisms by which the genes contribute to SRH have still to be determined. The single subjective item of SRH picks up the contributions from many background systems, including mental and physical health, as well as cognitive abilities and personality.

